# *De novo* designed protein enables precise epitope-level control of Gremlin-1 antagonism

**DOI:** 10.64898/2025.12.16.694677

**Authors:** Bunia I. Adsersen, Cecilie M. Møller-Jensen, Christian P. Jacobsen, Tanja Domeyer, Johannes R. Loeffler, Monica L. Fernandez-Quintero, Iacopo Tomberli, Astrid B. Jensen, Mikkel S. Jakobsen, Sandhya Bangaru, Andrew Ward, Esperanza Rivera-de-Torre, Stefán B. Gunnarsson, Andreas Hald, Timothy P. Jenkins

## Abstract

Growth-factor antagonists regulate key developmental and pathological processes, yet the molecular principles governing their control remain poorly understood. Gremlin-1 (GREM1) is a secreted antagonist of bone morphogenetic proteins (BMPs) whose dysregulation is implicated in fibrosis and cancer. Here, we report a *de novo* designed protein that binds GREM1 with sub-nanomolar affinity and selectively releases BMPs for downstream signalling. By integrating generative deep-learning–based protein design with molecular-dynamics–derived flexibility descriptors, we identify a predictive relationship between interface rigidity, desolvation energy, and binding success. The resulting binder, RF1-2, reproduces the native BMP-binding epitope on GREM1 at near-atomic precision, as confirmed by cryo-electron microscopy, and competitively blocks BMP-2 and BMP-4 association. These results establish interface rigidity as a key physical determinant of antagonist inhibition and demonstrate how AI-guided protein design can uncover molecular principles underlying extracellular signalling control.

## Introduction

Cells depend on growth factors for communication, differentiation, proliferation, and tissue regeneration in most organs [1–5]. The transforming growth factor *β* (TGF-*β*) super-family is among the most versatile of these regulatory systems, directing developmental patterning and maintaining tissue homeostasis throughout life [6–8]. Within this system, BMPs regulate processes ranging from skeletal formation to tumour suppression (Fig. 1a) [9–14]. Because these signals can drive both regeneration and pathological processes, BMP activity requires precise regulation. Excessive BMP signalling induces ectopic bone formation and abnormal differentiation, whereas insufficient signalling promotes fibrosis and malignant transformation [11, 15–17].

**Fig. 1:**
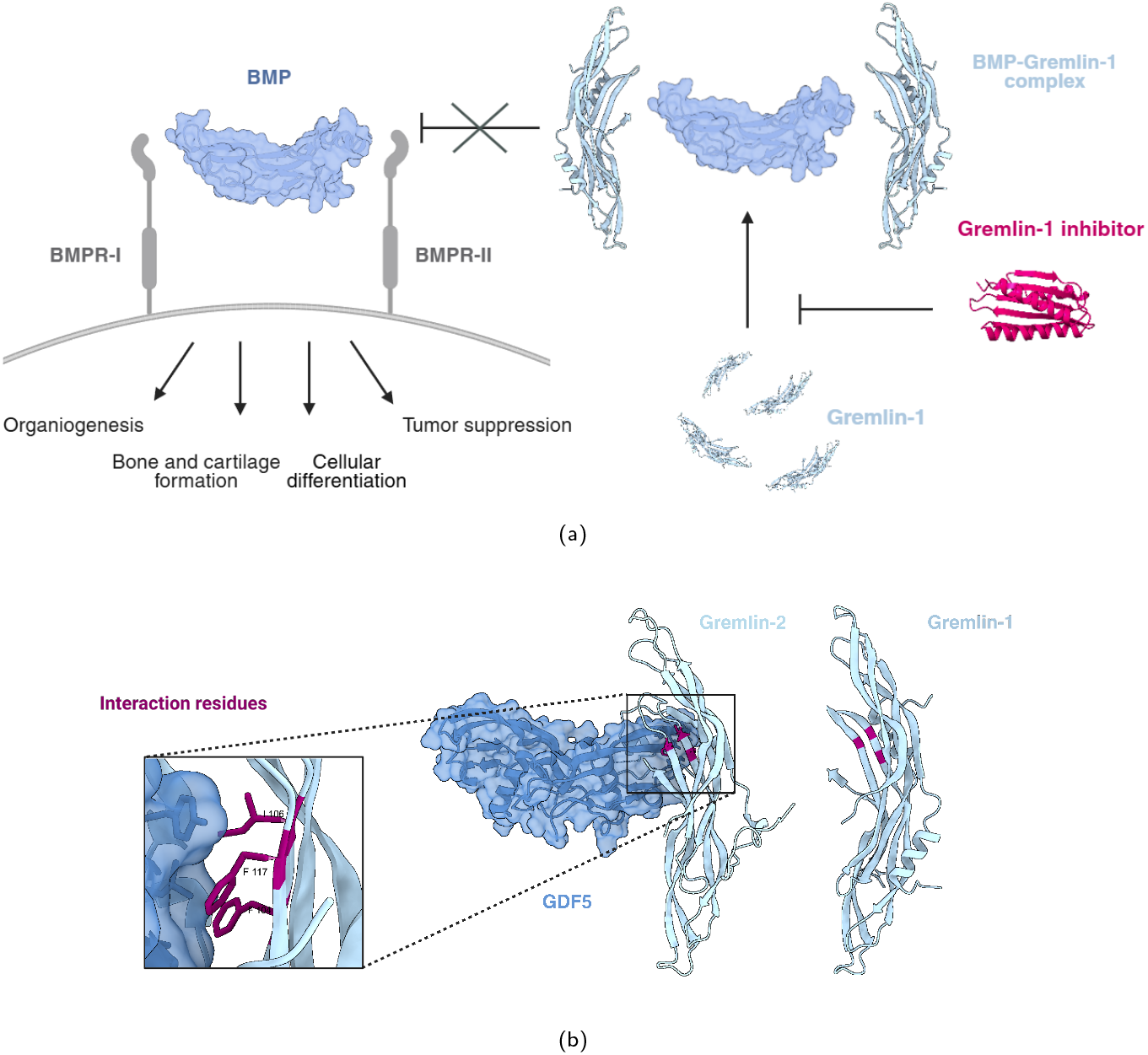
(a) Mechanism of GREM1 (PDB: 5AEJ) inhibition of BMP (PDB: 6OML) signalling and reversal by a GREM1 inhibitor, e.g. an anti-GREM1 designed miBd (AF2-*ig* prediction of RF1-2 miBd). (b) GREM1-BMP interaction residues inferred from GREM2-GDF5 complex (PDB: 5HK5).

This regulatory balance is maintained by a limited group of secreted antagonists that inhibit ligand–receptor binding. This includes GREM1, which serves as a principal antagonist of BMP-2, BMP-4, and BMP-7, functioning as a molecular brake on BMP signalling [18]. Dysregulated GREM1 expression disrupts the BMP pathways, and has been linked to pathological tissue remodelling associated with fibrosis, cardiovascular disease, and cancers [9, 19–21]. Despite these associations, the molecular mechanisms underlying GREM1 inhibition remain largely unresolved. We reasoned that designing a protein capable of binding GREM1 with atomic precision that neutralises a clinically relevant antagonist would enable us to elucidate the molecular principles governing growth-factor control.

To test this hypothesis, generative deep-learning methods for *de novo* protein design were applied to generate small protein binders, called minibinders (miBds), that engage GREM1 in the estimated GREM1–BMP interface. To this end, RFdiffusion and ProteinMPNN were used to generate diverse protein backbones and sequences, which were evaluated through co-folding with AlphaFold2-*initial guess* (AF2-*ig*) for structural compatibility [22–27]. Among 18 designs evaluated *in vitro*, four binders were identified, including one exhibiting sub-nanomolar affinity for GREM1 (approx. *K*_*d*_ = 0.3 nM). MD analyses revealed that true-positive binders possess a rigid, low-entropy interface with minimal desolvation costs. Furthermore, combining dynamic and static co-folding metrics enhanced the ability to distinguish true positives from false positives. Collectively, these findings highlight interfacial rigidity and low desolvation penalty as critical factors for high-affinity recognition, demonstrating that AI-guided protein design can reveal the physical principles underlying extracellular signalling.

## Results

### De novo design of GREM1 binders

The aim of the study was to generate a GREM1 binder that blocks the native ligand interaction with BMP-2 and BMP-4. However, the interaction between GREM1 and BMPs remains largely unexplored. To address this gap, we leveraged the high sequence homology between GREM1 and Gremlin-2 (GREM2; 71% sequence identity, excluding the less conserved N-terminal region) as both bind BMP-2 [28]. Notably, prior work has demonstrated that GREM2 leverages certain residues (F104, I106, F117) to bind both its high-affinity ligand BMP-2 and the lower-affinity ligand GDF5. These residues are structurally conserved in GREM1 (F125, I127, F138) suggesting a shared mechanism for BMP-2 recognition (Fig. 1b) [28, 29]. Accordingly, the GREM2-GDF5 complex (PDB: 5HK5) was used to infer key interacting residues between GREM1 and BMP-2.

A diverse range of miBds was generated against the GREM1 structure utilizing several design strategies. These included “unconditioned,” “hotspot” (targeting residues near the inferred GREM2-BMP-2 interaction site), and “*β*-sheet enriching” campaigns, resulting in 4,590 initial backbone designs. Partial diffusion was then applied iteratively to the top-scoring backbone designs, from each campaign, for further *in silico* refinement, yielding approximately 52,000 miBds. This iterative refinement continued until the improvement in the interface predicted alignment error (iPAE) metric plateaued (Supp. Fig. 5a). The resulting designs were filtered based on AF2-*ig* iPAE, binder predicted local distance difference test (pLDDT), and backbone diversity (Supp. Fig. 5b), leading to the selection of 18 miBds to undergo *in vitro* validation.

### In vitro validation of sub-nanomolar binding affinity and functional inhibition

Following the *in silico* design, all 18 selected miBds were expressed in a cell-free expression system, and four were identified to bind GREM1 by Luminescent Oxygen Channeling Immunoassay (LOCI). Three miBds (RF1-1, RF1-6, and RF2-2) demonstrated weak binding, whereas RF1-2 exhibited a strong interaction with GREM1 (Fig. 2a). Following the LOCI test, a full kinetic biolayer interferometry (BLI) experiment (Fig. 2d) demonstrated that RF1-2 exhibits sub-nanomolar binding affinity for GREM1 (approx. *K*_d_ = 0.3 nM, *R*^2^ = 0.99). A subsequent BLI experiment indicated that RF1-2 does not display non-specific binding to human insulin or SA biosensors.

**Fig. 2:**
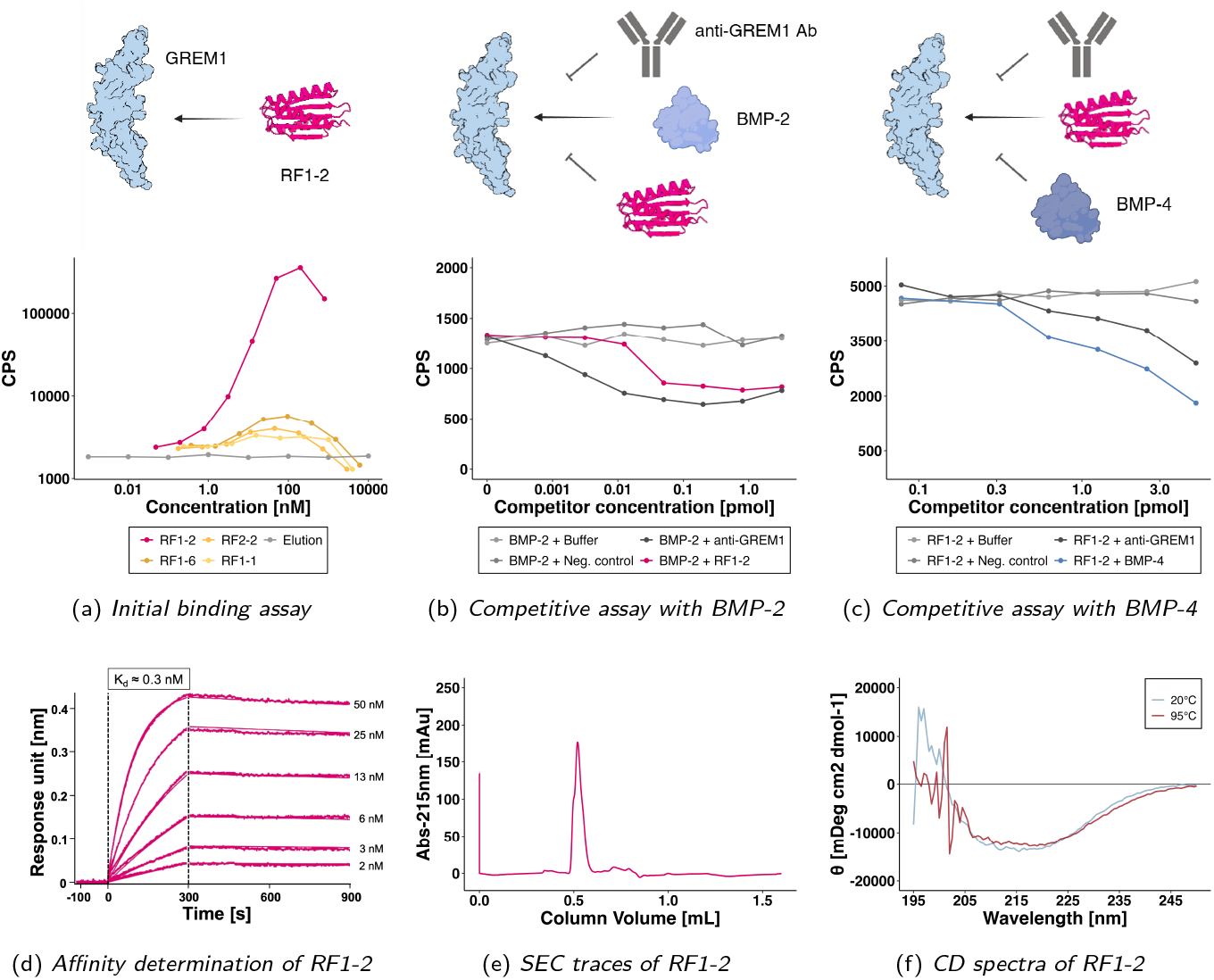
Functional assay and characterization results. (a) Initial binding assay to confirm GREM1 binding potential. (CPS: counts per second). (b) Competitive inhibition of BMP-2. Constant concentration of BMP-2 and GREM1. (anti-GREM1: confirmed competitor of BMP-2. CPS: counts per second). (c) Competitive inhibition of BMP-4. Constant concentration of RF1-2 and GREM1. (anti-GREM1: confirmed competitor of BMP-4. CPS: counts per second). (d) BLI sensograms of RF1-2 binding to immobilized Gremlin-1 for affinity determination. (e) SEC traces of purified RF1-2. (mAu: milli-absorbance units). (f) CD profiling of RF1-2 confirming *αβ*-secondary structure and thermal stability. (*θ*: molar ellipticity).

With high-affinity and specific binding to GREM1 established, we next used LOCI to test whether RF1-2 could outcompete BMP-2 and BMP-4 in binding to GREM1. In the competitive BMP-4 experiment, GREM1 was used at 5,000 pM with a BMP-4 dilution series from 500 pM, and RF1-2 at 2 nM. Under these conditions, RF1-2 produced a clear, concentration-dependent inhibition of BMP-4 binding, with no signal from the negative control or buffer (Fig. 2c). In a second competitive assay using BMP-2, GREM1 was present at 40 nM, BMP-2 at 20 nM, with a RF1-2 dilution series from 320 nM. RF1-2 again demonstrated concentration-dependent competitive inhibition, confirming its ability to disrupt BMP-2 and BMP-4 interactions with GREM1. RF1-2 achieved BMP blocking comparable to anti-GREM1 antibodies (Fig. 2b, 2c). Furthermore, in a competitive LOCI assay with an in-house developed antibody (anti-GREM1 [30]), RF1-2 exhibited concentration-dependent competitive binding (Supp. Fig. 6). Following binding characterization, RF1-2’s aggregation propensity was assessed via size-exclusion chromatography (SEC). The SEC profile demonstrated that the purified miBd was monodisperse, eluting as a single peak at the expected monomeric molecular weight based on the column calibration curve (Fig. 2e); thus indicating no significant aggregation or dimerization. Circular dichroism (CD) spectroscopy further confirmed that RF1-2 adopts the expected fold, with a spectrum characteristic of both *α*-helical and *β*-sheet secondary structures and exhibited excellent thermostability, with no indication of denaturing even at 95°C (Fig. 2f).

### Using molecular dynamics for retroactive improvement of experiment precision

Although the initial design campaign yielded a picomolar GREM1 binder (RF1-2) on the first attempt, several other designs with similarly high AlphaFold-derived confidence scores did not exhibit detectable binding *in vitro*. This outcome suggests that static co-folding metrics, including interface pLDDT, ipTM, and iPAE, which are foundational to recent improvements in *de novo* binder design success rates [22, 23, 27], may fail to capture the essential energetic and dynamic features of protein–protein interactions.

To determine whether incorporating protein dynamics could reduce false positives, we retrospectively computed molecular dynamics (MD)-derived physicochemical and dynamic descriptors for each design. We then evaluated their ability to improve discrimination of binders identified in the LOCI experiment (Fig. 2a), in which 4 of 18 designs bound GREM1. For each of the 18 miBds, we conducted four independent 500-nanosecond simulations, totalling approximately 2.0 microseconds per binder. From these trajectories, we quantified recurrent interface contacts (present in at least 10% of simulation time) and derived dynamic descriptors that capture both local and global flexibility, as well as electrostatic contributions to the binding interface. Local interface flexibility was assessed using per-residue B-factors (derived from root mean square fluctuation, RMSF) and a *χ*-entropy metric reflecting side-chain dihedral entropy. As several interfaces were highly charged, with up to 35 salt bridges, we estimated desolvation penalties to account for the energetic cost of burying charged residues. Finally, overall complex dynamics were summarized by clustering the MD trajectories using hierarchical clustering to obtain a cluster count descriptor (Fig. 3a). When we assessed correlations between MD-derived metrics and static co-folding metrics, iPAE from AF-2 showed higher absolute Spearman correlations with dynamic features, including number of clusters (*ρ* = 0.60), *χ*-entropy (*ρ* = 0.41), and interface B-factors (*ρ* = 0.32), compared to AlphaFold-3 (AF3) ipTM. In contrast, AF3 ipTM exhibited a higher absolute correlation with the desolvation penalty (*ρ* = − 0.21; Supp. Fig. 7). Notably, AF3 ipTM also correctly identified the picomolar binder as having the highest ipTM score among the 18 tested designs.

**Fig. 3:**
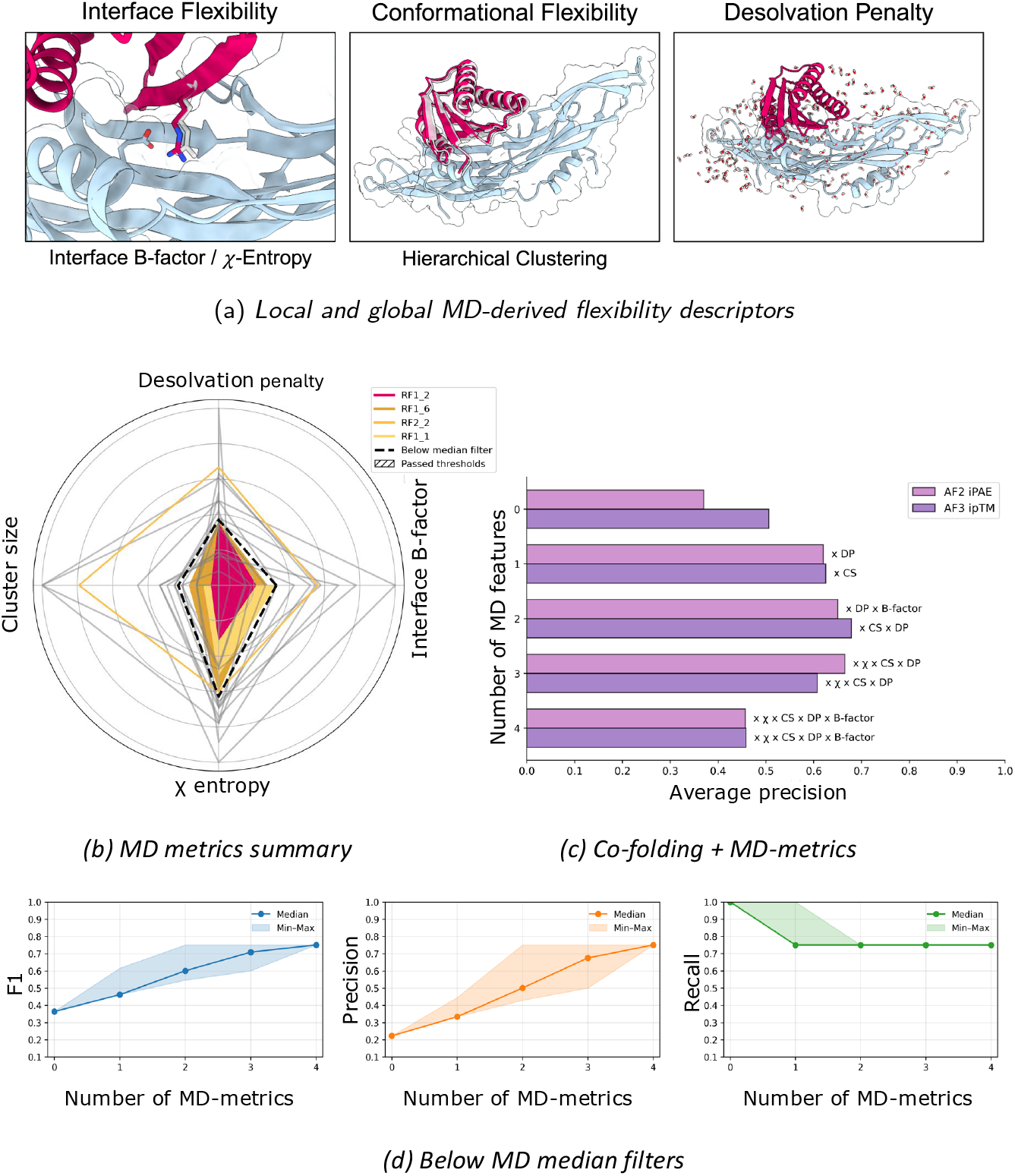
MD descriptors. (a) Schematic of what aspect of interface dynamics each MD-derived descriptor captures. (b) Distributions of the four MD metrics across the 18 miBds tested *in vitro*; the black dotted line marks the per-metric median, and filled squares indicate designs passing all four below-median filters. (c) Average precision (AP) when augmenting static co-folding metrics with MD descriptors; for each feature count, the best-performing combination is shown. (DP: Desolvation penalty, CS: Cluster size) (d) F1, precision, and recall using simple below-median decision cutoffs to classify binders in LOCI; lines denote medians and shaded bands show the observed minima and maxima.

Using these descriptors, we evaluated whether augmenting static co-folding metrics with MD-derived features improves discriminative performance in identifying true binders from false positives. To this end, we computed precision–recall (PR) curves and summarized performance as average precision (AP) across pairwise and multi-feature combinations of the static co-folding metrics and MD descriptors. This analysis was performed for both iPAE from the AF2-*ig* model used during candidate selection, and the ipTM score from AF3 [31]. Using iPAE (AF2-*ig*) combined with a single MD descriptor (iPAE x desolvation penalty) increased the AP by up to ~ 25%, compared to iPAE as a baseline. Adding one additional descriptor (iPAE x desolvation penalty x number of cluster) yielded a further ~ 4% improvement (Fig. 3c). Repeating this analysis with AF3 ipTM showed similar trends: ipTM combined with the number of clusters improved AP by ~ 12%, and adding the desolvation penalty provided an additional ~ 5% increase. In both cases, larger feature sets did not help and often reduced performance, indicating diminishing returns beyond two features. We also investigated whether relying solely on MD-derived metrics could improve performance by applying a simple below-median threshold rule to each MD descriptor (Fig. 3b). Using this rule across all MD descriptors retained three designs, all of which were binders in the LOCI assay, while excluding all non-binders. Notably, RF1-2 (sub-nanomolar affinity) exhibited the lowest local and global flexibility among the downselected designs, reflected by the lowest *χ*-entropy, lowest interface B-factor, and smallest number of clusters. To systematically compare MD features, we computed precision, recall, and F1 scores across all feature combinations using below-median decision cutoffs (Fig. 3d). Relative to the AF2-ig iPAE/pLDDT baseline, which effectively labelled all 18 designs as binders, a single MD feature improved the median F1 by +9.8% (best descriptor: *χ*-entropy). Combining two features increased the median F1 by ~ 23.6% (best descriptor pair: desolvation penalty + *χ*-entropy), with performance plateauing for larger feature sets.

### Experimental confirmation of epitope control

To confirm that the designed binder engages the intended BMP-binding epitope, we determined the structure of the GREM1–RF1-2 complex by cryo-electron microscopy (cryo-EM). The resulting model-to-map fit supports the predicted binding pose, with RF1-2 adopting a fold and interface orientation consistent with the AF2-*ig* design model.

Despite the modest overall resolution ( ~ 5 Å), the density clearly defines the relative placement of the binder and target, positioning RF1-2 across the BMP-contacting surface of GREM1. Orientation refinement, achieved by maximising the number of atoms within the cryo-EM density, resulted in *>*90% of model atoms contained within the map, indicating a robust fit between prediction and experiment.

The design targeted residues F125, I127, and F138, key determinants of BMP recognition, and the cryo-EM density is consistent with engagement of this hydrophobic patch. MD simulations further support this epitope assignment, revealing that F125 and F138 form recurrent contacts with RF1-2 throughout trajectories, whereas I127 remains buried and solvent-inaccessible, as expected for a core interface residue. Together, these structural and dynamic data confirm that RF1-2 binds precisely at the intended BMP-binding surface of GREM1, validating the design hypothesis and providing a structural rationale for its inhibitory activity.

**Fig. 4:**
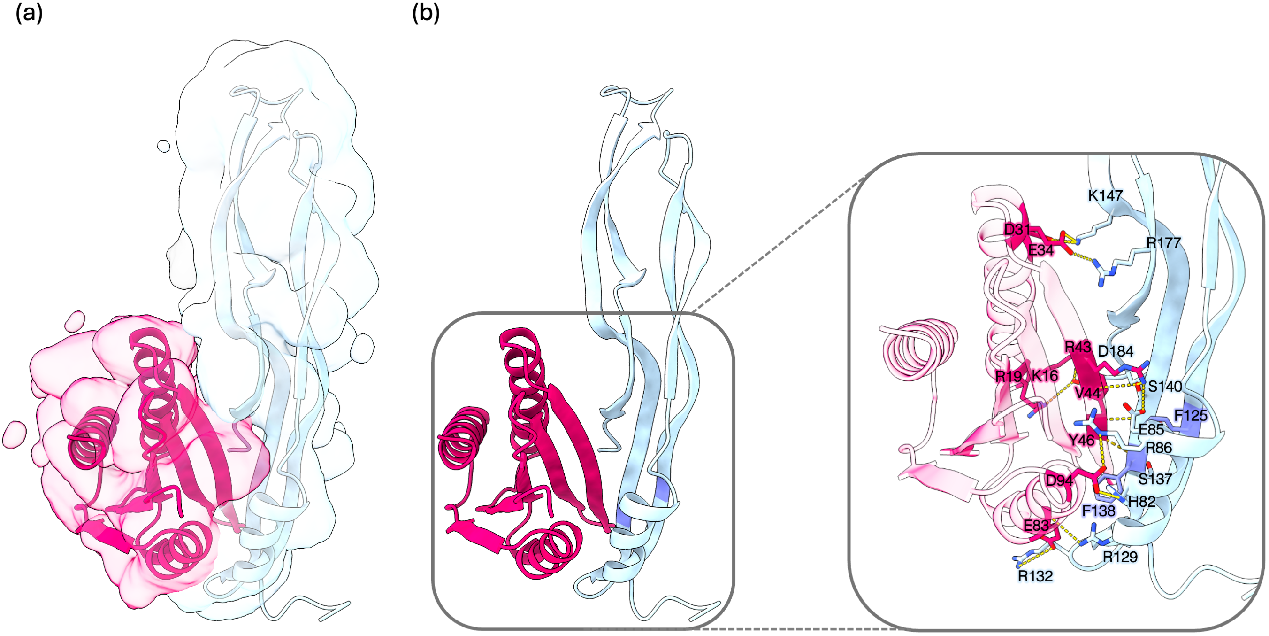
(a) Cryo-EM density map (4.8 Å) of the GREM1 (blue) in complex with RF1-2 (pink) indicating a binding pose consistent with the predicted docked in AF3 structure model. (b) Hydrogen bond and Van der Waals interactions of RF1-2 with GREM1 characterized by MD simulations, highlighting residues F125 and F138 in dark blue.

## Dicussion

Growth factors play a central role in balancing cell proliferation and differentiation, governing critical processes such as wound healing, tissue regeneration, and embryonic development [4]. When these signalling pathways become dysregulated, they can drive fibrosis and cancer, highlighting the need for molecular tools capable of dissecting and precisely modulating growth-factor signalling [5, 8]. Significant progress in protein structure prediction and computational design is now making it possible to generate functional binders with greater speed and precision, while being cost-efficient. Here, we demonstrate that generative *de novo* design rapidly can yield functional binders for signalling proteins. Within just three weeks, an unconditional RFdiffusion campaign, combined with ProteinMPNN and AF2-*ig* validation, created a binder with sub-nanomolar affinity that effectively inhibits the BMP antagonist GREM1.

Of 18 candidates tested *in vitro*, four were identified as binders and one, RF1-2, was found to bind GREM1 at nanomolar concentrations in both BLI and LOCI assays. RF1-2 targets a conserved hydrophobic patch (F125, I127, F138) on GREM1, corresponding to an inferred BMP-binding epitope deduced from the GREM2–GDF5 structure, and cryo-EM confirmed this interface despite moderate map resolution ( ~ 5 Å). The binder also demonstrated a melting temperature above 95°C and no evidence of aggregation, supporting the translational potential of AI-guided scaffolds.

Notably, RF1-2 emerged from a fully unconditional generative run with no explicit epitope constraints. This suggests that granting diffusion models greater backbone freedom may yield scaffolds intrinsically compatible with natural protein–protein geometries, whereas rigid hotspot specifications could bias designs away from productive interfaces. Thus, the success of RF1-2 reflects not only the ability of *de novo* design models to accurately model protein–protein interactions, but also the implicit physical priors it has learned. Our retrospective molecular-dynamics (MD) analysis clarified why RF1-2 succeeded where others failed, revealing that true-positive binders share rigid, low-entropy interfaces with favourable desolvation energetics, features absent in nonbinders. This analysis further revealed that augmenting static AlphaFold confidence metrics with dynamic and energetic descriptors increased predictive precision by up to 25% (AF2-*ig*) and 12% (AF3), indicating that integrating larger-scale dynamics simulations into protein–protein interaction prediction improves performance. Incorporating such dynamic constraints into future generative workflows might markedly reduce false positives and improve design reliability.

In summary, generative protein design enables the zero-shot creation of high-affinity protein binders, and augmenting co-folding predictions with physics-based scores can improve discrimination between true binders and false positives. Together, these elements outline a generalizable and efficient path toward creating research- and potentially therapeutic protein binders that can probe and manipulate growth factor signalling in health and disease.

### Limitations of the study

This proof-of-principle study remains limited by the small number of experimentally validated designs, and future benchmarking across additional targets will be essential to generalise the MD-derived predictors. Moreover, while the rigid *α*-helical bundle architecture confers exceptional stability and affinity, it may restrict access to concave or recessed epitopes, motivating exploration of hybrid scaffolds that balance rigidity with conformational flexibility. Even within these bounds, the present results demonstrate that combining generative models with physics-based refinement can deliver functional, high-affinity binders to challenging targets within weeks.

## Methods

### De novo design of miBds with RFdiffusion

The computational design of binding proteins was performed against the homodimeric formation of GREM1, as this is the biological assembly of the protein [9]. Chain A (residues 71-184) and chain B (residues 72-184) from the GREM1 structure (PDB: 5AEJ), were selected as target for RFdiffusion [22]. Binders were designed within the range of 100-150 amino acids and several campaigns were initiated to generate a diverse range of binders including unconditioned, hotspot, and *β*-sheet enriching campaigns. Sequences were generated using ProteinMPNN, with Vanilla weights and a sampling temperature of 0.1, followed by validation with AF2-*ig* [24, 25, 32]. The best scoring initial backbone structures underwent partial diffusion for further *in silico* refinement. The degree of noise (“diffuser.partial_T”) added to the original structures were varied with input-values 15, 20, and 25. The binder designs were filtered based on two main AF2-*ig* metrics, iPAE *<* 10 and binder pLDDT *>* 85. All best scoring structures were visually inspected to assess binder orientation, availability of C-terminal for tags included in expression and assays, and overall structural credibility. Furthermore, the AF2-*ig* predicted GREM1 binding epitopes for each binder were examined for overlap with the predicted BMP-2 binding residues (F125, I127, F138) [29].

### Expression

#### Cell-free miBd expression

All miBds were expressed in a cell-free system from human-codon–optimized DNA fragments reconstituted in milliQ water. PUREfrex 2.1 components Solution I, cysteine, and GSH were thawed at room temperature for one minute to completely dissolve and then cooled on ice. Solution II, and III were thawed on ice. Each solution was mixed by vortex and centrifuged briefly, whereafter they were combined to a mastermix. Then, 24µL mastermix and 1 µL of DNA were added per well in a 96-well plate and incubated overnight at 37°C with shaking. After incubation, 15 µL of pre-washed Pierce™ Ni-IMAC MagBeads per well was used to capture His-tagged proteins at room temperature (RT) for 60 min, followed by PBS washing and elution with 100 µL of 250 mM imidazole for 30 min. Protein expression was confirmed by SDS-PAGE. Mass and concentration were determined by reverse-phase LC-MS.

#### Recombinant expression of RF1-2

The miBd RF1-2 was expressed in E. coli using a commercially synthesized plasmid (pET42c(+), GenScript). To enhance solubility and facilitate purification, the fusion protein contained, from the N-terminus: a glutathione S-transferase (GST) tag, followed by a 6×His tag, an S-tag, and RF1-2, each separated by linker sequences.

RF1-2 was expressed in E. coli BL21(DE3) cells transformed with the plasmid via heat-shock transformation. Transformed cells were selected on LB agar plates supplemented with 50 µg/mL kanamycin and grown overnight at 37 °C. A single colony was used to inoculate a 10 mL LB preculture containing kanamycin, incubated overnight at 37 °C with shaking (220 rpm). For expression, cells were inoculated (1:100) into ZY based autoinduction medium (ZY medium base supplemented with 2 mM *MgSO*_4_, 1% 50×5052, 1% 50× M, 0.02% 1000×metals mix, and 50 µg/mL kanamycin), and incubated overnight at 37 °C, 220 rpm. Cells were harvested by centrifugation (6,000×g, 20 min), resuspended in 50 mM Tris, 200 mM *NaCl*, 20 mM imidazole, 0.1% Tween-20, pH 8.0, and lysed by sonication (40% amplitude for 7 min) on ice. The lysate was clarified by centrifugation (17,000×g, 30 min, 4 °C), and the soluble fraction was purified using a Cytiva HisTrap™ 5 mL column on a Bio-Rad NGC FPLC system. The column was equilibrated with 50 mM Tris, 200 mM NaCl, 20 mM imidazole, pH 8.0 and bound proteins were eluted with a linear gradient up to 250 mM imidazole. Fractions were analysed by SDS-PAGE. GST was removed by thrombin digestion (2 Units per 1 mg protein) following buffer exchange into thrombin-compatible buffer (50 mM Tris, 200 mM *NaCl*, 2.5 mM *CaCl*_2_, pH 8.0). Cleaved protein was separated from the GST tag using *Ni*^2+^ affinity chromatography with HisPur™ resin in a gravity-flow format. Protein purity was monitored by SDS-PAGE at each step.

### Luminescent Oxygen Channeling Immunoassay

Luminescent Oxygen Channeling Immunoassay (LOCI) / AlphaLISA is a bead-based immunoassay method. Binding interactions bring Streptavidin-coated Donor Beads (6760002L, Revvity) and Acceptor Beads (6772004B, Revvity) close together. Excitation of the donor beads upon illumination at 680 nm causes the formation of singlet oxygen, which is transferred to the acceptor beads in close proximity, resulting in measurable luminescence[33]. In these assays, Streptavidin-coated donor beads bind to a biotinylated molecule and acceptor beads are conjugated with anti-His-tag antibodies or anti-GREM1 antibodies.

#### Initial binding assay

For the experiment, 20 µL of each miBd sample was serially diluted 4-fold in LOCI buffer across eight concentrations in a PCR plate. Negative controls (LOCI buffer and IMAC elution buffer) were prepared in separate columns. Then, 1 µL of each sample was transferred to a 384-well plate in duplicates. Acceptor beads conjugated with anti-His-tag antibodies were diluted 1:150 in LOCI buffer, and biotinylated GREM1 was diluted to 8 nM in PBS containing 1% BSA and 0.05% Tween. 15 µL of this mixture was added to each well using a Tempest^®^ Liquid Handler, followed by a 60 min incubation. Donor beads were diluted in LOCI buffer, and 30 µL was added to each well, followed by an additional 30 min incubation. The plate was read using a PerkinElmer EnVision™ 2102 Multilabel Reader to measure luminescence.

#### Competitive inhibition of anti-GREM1 antibody

GREM1 was diluted in PBS with 1% BSA and 0.05% Tween20 to a final concentration of 2 nM. RF1-2 was prepared in a 14 sample serial dilution with a dilution factor of 3 and a maximal concentration of 800 nM. 1 µL of GREM1 and RF1-2 was added to a 384-well AlphaPlate alongside 8 µL LOCI buffer, and the plate was incubated for 60 min. Biotinylated competing anti-Gremlin antibody (Novo Nordisk) with a concentration of 0.5 nM was mixed with 10 µL acceptor beads conjugated with a non-competing anti-Gremlin antibody (Novo Nordisk). 20 µL of the mix was added to each well and the plate was incubated for 10 min. 20 µL donor beads diluted in LOCI buffer was added to the wells and the plate was incubated for 10 min before reading the luminescence on a PerkinElmer EnVision™ 2102 Multilabel Reader.

#### Competitive inhibition of BMP-4 binding to GREM1

GREM1 was diluted to 5000 pM in PBS with 1% BSA and 0.05% Tween20. BMP-4 (ab231930, Abcam) was diluted in LOCI buffer to prepare concentrations of 500, 250, 125, 63, 31, 16, 8 and 0 pM. A competing anti-GREM1 antibody was diluted to the same concentrations to be used as a positive control. The non-competing antibody and buffer were used as negative controls. 10 µl of BMP-4, competing and non-competing antibody dilutions, as well as buffer, were added to a 384-well AlphaPlate in duplicates. 1 µl of GREM1 was added to all wells and the plate was incubated for 30 min. 15 µl of 33.33 µg/ml AlphaLISA acceptor beads conjugated to an anti-His-tag antibody (MA1-21315-1MG, Invitrogen), 8 nM of another biotinylated non-competing anti-GREM1 antibody (Novo Nordisk) and 2 nM miBd RF1-2 were added to each well. After 60 min incubation at RT, 20 µl of 100 µg/ml Streptavidin-coated donor beads were added to each well and the plate was further incubated for 30 min before reading the luminescence on a PerkinElmer EnVision™ 2102 Multilabel Reader.

#### Competitive inhibition of BMP-2 binding to GREM1

GREM1 was diluted to 40 nM in PBS with 1% BSA and 0.05% Tween20. Biotinylated BMP-2 (bs-48015P-Biotin, Bioss) was diluted in LOCI buffer to a concentration of 20 nM. RF1-2 and the competing antibody as positive control, and a non-competing antibody as a negative control were prepared in a 7 sample serial dilution with LOCI buffer with a dilution factor of 4 and the maximal concentration of 320 nM. 1µL of GREM1 was transferred to a 384-assay plate with 3 µL of LOCI buffer. 10µL of biotinylated BMP-2, RF1-2, antibodies, and LOCI buffer were added to their respective columns of the assay plate. 15 µL of 33.33 µg/ml AlphaLISA acceptor beads conjugated with another non-competing antibody were added and the plate was sealed and incubated over night at RT. 15 µL of 133.34 µg/ml Streptavidin-coated donor beads were added to each well and the plate was incubated for 30 min before reading the luminescence on a PerkinElmer EnVision™ 2102 Multilabel Reader. The final molar quantity of GREM1 and BMP-2 in each well were 40 fmol and 200 fmol, respectively.

### Biolayer Interferometry

Biolayer interferometry (BLI) experiments were performed on an Octet® RED384 system (Sartorius) using streptavidin (SA) biosensors (Sartorius item no. 18-5019). Biosensor were pre-equilibrated in kinetic buffer prior to use. All ligand and analyte dilutions were prepared in kinetic buffer. The kinetic buffer consisted of 20 mM HEPES supplemented with 1 mg/ml IgG-free bovine serum albumin, 0.05% v/v Tween 20, and 5 mM CaCl_2_, adjusted to pH 7.4. The regeneration buffer consisted of 10 mM glycine-HCl, adjusted to pH 1.5. Experiments were run at 21°C with shaking at 1000 rpm, and data were processed with Data Analysis HT software (Sartorius) v.12.0.2.59.

#### Characterization of the binding kinetics of RF1-2 for human GREM1

For determination of binding kinetics, SA biosensors were initially conditioned by three cycles of incubation in regeneration buffer for 5 sec, followed by incubation in kinetic buffer for 5 sec. The conditioned SA biosensors were then sequentially incubated in kinetic buffer for 30 sec, biotinylated human GREM1 (ligand) for 300 sec, kinetic buffer for 120 sec, RF1-2 (analyte) for 300 sec, and kinetic buffer for 600 sec. Biotinylated human GREM1 was immobilized on the SA biosensors at ligand concentration of 0.8 *µ*g/mL. The binding of RF1-2 to the ligands was tested at six different analyte concentrations (50, 25, 12.5, 6.25, 3.13, 1.56 nM). For data correction, SA biosensor H served as a reference sensor without analyte (RF1-2) and was used to subtract baseline drift from the other SA biosensors. The association (*k*_on_) and dissociation (*k*_off_) rate constants were determined by globally fitting the sensorgrams to a 1:1 binding model. The equilibrium dissociation constant (*K*_d_) was calculated as (*k*_off_)/(*k*_on_).

#### Characterization of RF1-2 for Non-Specific Binding to human Insulin

For characterisation of non-specific binding, SA biosensors were initially conditioned by three cycles of incubation in regeneration buffer for 5 sec, followed by incubation in kinetic buffer for 5 sec. The conditioned SA biosensors were incubated sequentially in kinetic buffer for 30 sec, followed by biotinylated human insulin (ligand) or biotinylated human GREM1 (ligand) for 300 sec, kinetic buffer for 120 sec, RF1-2 (analyte) for 300 sec, and kinetic buffer for 300 sec. Biotinylated human insulin or biotinylated human GREM1 was immobilized on the SA biosensors at a ligand concentration of 10 *µ*g/mL. The binding of RF1-2 to immobilized ligands and SA biosensor was tested at seven different analyte concentrations (50, 25, 12.5, 6.25, 3.13, 1.56 and 0,781 nM). For data correction, SA biosensor H served as a reference sensor without analyte (RF1-2) and was used to subtract baseline drift from the other SA biosensors.

### Analytical Size Exclusion Chromatography

Analytical size-exclusion chromatography (SEC) was performed using a Bio-Rad NGC FPLC system to evaluate the monodispersity of purified RF1-2. A Superdex 75 10/300 GL column (Cytiva) was equilibrated with one column volume ( ~ 24 mL) of buffer (50 mM Tris, 200 mM *NaCl*, 2.5 mM *CaCl*_2_, pH 8.0). A 500 µL protein sample (concentration: 19.8 µM) was injected using a sample loop and eluted with 1.2 column volumes of the same buffer. Protein elution was monitored at 215 nm to assess the oligomeric state and potential aggregation.

### Circular Dichroism Spectroscopy

Far-UV circular dichroism (CD) spectroscopy was performed to assess the secondary structure and thermal stability of the purified RF1-2 at a concentration of 9.9 µM. CD spectra were acquired using a JASCO J-1500 spectropolarimeter (Easton, MD, USA) with a 0.1 mm pathlength quartz cuvette. Spectra were recorded at 20°C and 95°C to evaluate the thermal-induced unfolding, by averaging three scans in the 190–260 nm range with a 0.5 nm step size. The spectra were processed using JASCO SpectraManager software.

### Molecular dynamics

To test if MD could better discriminate true from false protein binders, we developed a protocol that applied MD simulations to the 18 selected miBds that had undergone *in vitro* validation. For each miBd, we conducted four independent MD simulations, each spanning 500 ns, resulting in an aggregate simulation time of 2 µs per binder. From these simulations, we extracted key interaction features at the interface, including hydrogen bonds and salt bridges, annotated with their respective contact frequencies. Interface residues were defined as those involved in non-covalent interactions for at least 10% of the total simulation time, providing a dynamic characterization of the binding interface. To assess the conformational stability of these interface residues, we calculated a per-residue interface B-factor, serving as a proxy for local flexibility. High B-factors at the interface may indicate entropically unfavorable regions that could disrupt stable binding. Additionally, since several miBds featured highly charged interfaces, some forming up to 35 salt bridges, we incorporated an estimate of the desolvation penalty to account for the energetic cost of burying charged residues. To do this, we applied residue-specific desolvation penalties derived from empirical data on PPI interfaces. This allows us to down-select binders with unrealistically high electrostatic costs, which are unlikely to form stable complexes under physiological conditions.

To assess whether MD-derived descriptors could improve the discriminative power of static co-folding metrics, we min–max normalized the four MD features and the co-folding confidence scores to [0, 1] and added a small offset (*ε*) to avoid zeros. To harmonize directions, we inverted the MD features and iPAE (lower-is-better) as (*x*^*′*^ = 1 − *x*), aligning them with ipTM so that larger values indicate more favorable predictions across all features. For each baseline (reoriented AF2-*ig* iPAE or native AF3 ipTM), we then constructed composite interaction scores by element-wise multiplication with every combination of MD features ((*k* = 1 … 4)). For each composite score, we computed precision–recall (PR) curves and summarized performance as average precision (AP) using LOCI-derived binder/non-binder labels.

Secondly, to evaluate discriminative performance of simple decision cut-off rules, below-median threshold scores were calculated per feature. For combinations of (*k*) features ((*k* = 1 … 4)), a design was classified as a predicted binder only if it passed all (*k*) thresholds. Using the LOCI labels, we then computed precision, recall, and F1 scores for each combination and summarized performance across (*k*).

### Cryo-EM structural determination of GREM1 - RF1-2 complex

For cryo-EM sample preparation, recombinantly expressed GREM1 was cleaved using Factor Xa (NEB, Cat #P8010S) to remove the C-terminus using the manufacturer’s protocol. Briefly, 1 mg of GREM1 was incubated with 10 µg of Factor Xa and incubated for 5 hrs at RT prior to the addition of RF1-2 (2 M excess to GREM1). The complex was further incubated for 60 min and purified over a Superose 200 increase 10/300 column (Cytiva). The fraction corresponding to the GREM1-RF1-2 complex was used for cryo-EM sample preparation. The final concentration of the complex on the grids was 0.5 mg/mL. The samples were added to glow discharged 1.2/1.3 Au Ultrafoil 300 mesh grids and subjected to vitrification using a Vitrobot Mark IV system. The settings were as follows: 3 µL of sample, temperature inside the chamber was 4°C, humidity was 100%, blotting force was 1, and wait time was 4 s. The sample was blotted off for 4.5 s and the grids were plunge-frozen into liquid-nitrogen-cooled liquid ethane.

Cryo grids of the complexes were imaged at 190,000× nominal magnification using a Falcon 4i camera on a Glacios microscope at 200 kV. Automated image collection was performed using EPU from ThermoFisher. Images were aligned, dose-weighted, and Contrast Transfer Function (CTF)-corrected in the CryoSPARC Live™ software platform, with automated image collection also performed using Smart EPU software (ThermoFisher). Data processing for all was carried out in CryoSPARC v4.7.0 [34]. Blob particle picking was performed on all micrographs with a minimum particle diameter of 80 Å and a maximum of 120 Å. Particles extracted at 304 pixels box size were used to perform multiple rounds of 2D classification, which were then used to generate a 3D reference model from abinitio refinement, followed by multiple rounds of hetero- and homogenous refinements to obtain good classes for further non-uniform and local refinements. Gold-Standard Fourier Shell Correlation resolution was calculated to be 4.76 Å. We docked the models into the cryo-EM density map in UCSF ChimeraX [35].

## Resource availability

### Lead contact

Requests for further information and resources should be directed to the lead contact, Timothy P. Jenkins.

### Materials availability

Materials can be made available upon reasonable request.

### Data and code availability

Code explanation and examples of binder design using RFdiffusion can be found at https://github.com/RosettaCommons/RFdiffusion#binder-design. The EM map of RF1-2 in complex with GREM1 has been deposited at the Electron Microscopy Data Bank (EMDB) with the accession code EMD-74525.

## Acknowledgements

We acknowledge the teams, institutions and grants that support open access databases and code, including AlphaFold2 and RFdiffusion, which were essential to this work. We thank members from the Center for Translational Protein Design for helpful discussions and input regarding both design strategies and laboratory validations. This work was supported by funds provided by Novo Nordisk A/S.

## Author contributions

T.P.J. and A.H. conceptualized the project. T.P.J, A.H., C.M.M, B.I.A., and C.P.J. administrated the work. B.I.A. and C.M.M. conducted the computational design and analyses of relevant GREM1 binding proteins. A.B.J. conducted cell free expression and purification of binding proteins. T.D. and S.B.G. designed and performed initial LOCI of binding protein abilities with assistance from B.I.A. and C.M.M.. M.S.J. conducted affinity binding measurements using BLI. T.D. designed and performed competing functional assays with IgG, BMP-2, and BMP-4 as the competitors of the designed binding proteins. E.R. and I.T. recombinantly expressed and purified binding proteins and performed SEC and CD to evaluate protein characteristics. J.L. and M.F. performed molecular dynamic predictions of relevant binding proteins. C.P.J. conducted performance evaluation of molecular dynamics descriptors. S.B. and M.F. performed preparatory experiments before M.F., S.B., and A.B.W. conducted cryo-EM structure determination of the GREM1 - RF1-2 complex. T.P.J., A.H., and C.P.J. provided research support and supervision.

## Declaration of generative AI and AI-assisted technologies in the writing process

During the preparation of writing this work the authors did not use AI or AI-assisted technologies.

## Supplementary Information

**Fig. 5:**
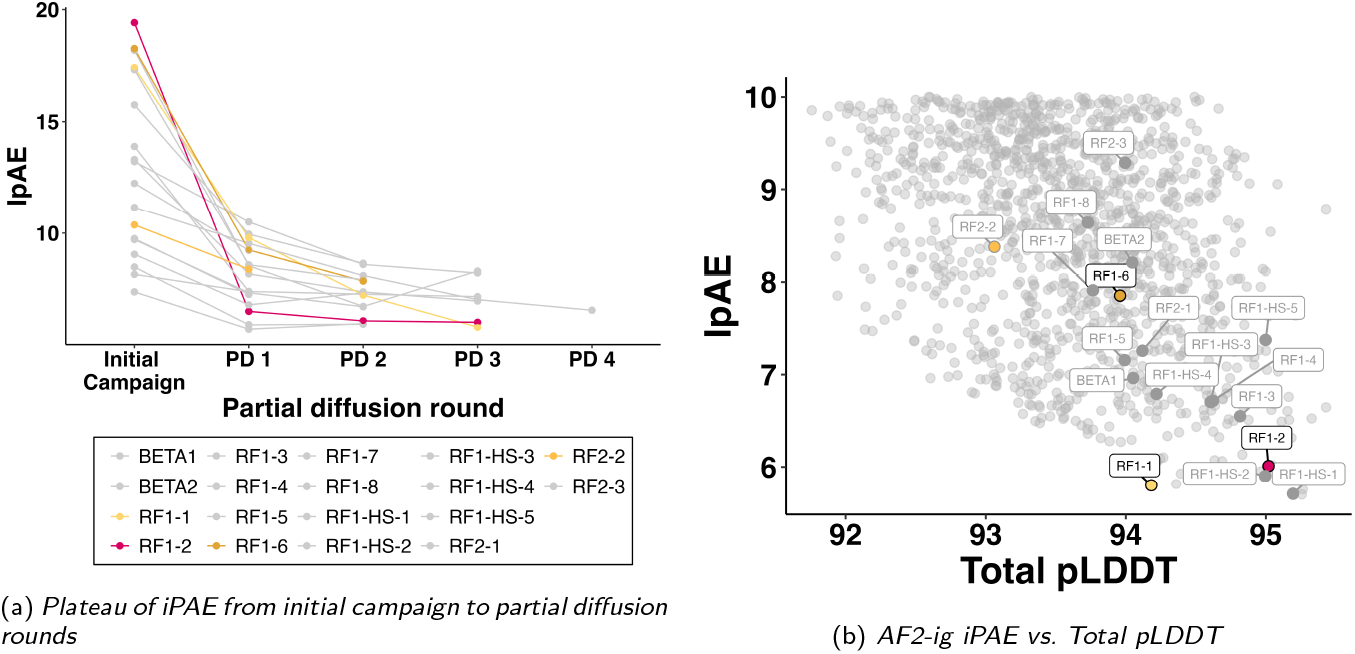
(a) AF2-*ig* iPAE of initial 18 miBd designs followed by their respective decline in iPAE during rounds of partial diffusion. (b) AF2-*ig* iPAE and total pLDDT of all miBd designs, zoomed in on the final 18 miBds chosen for *in vitro* validation. The highlighted binders are the ones filtered out by the molecular dynamics analysis as high potential binders.

**Fig. 6:**
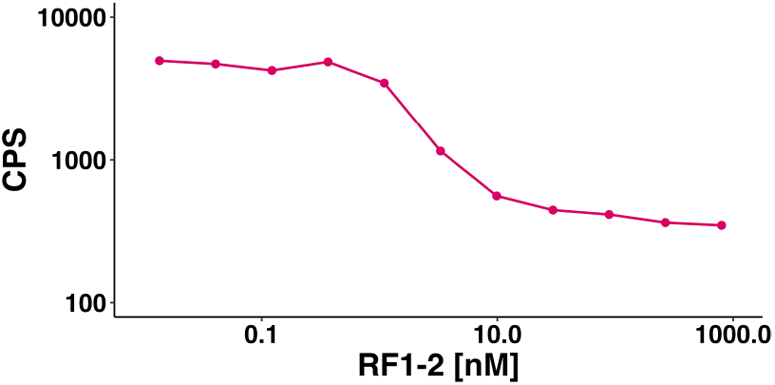
Concentration dependent competitive inhibition of in-house anti-GREM1 antibody (anti-GREM1). Constant concentration of GREM1 and biotinylated anti-GREM1, while RF1-2 concentration varies.

**Fig. 7:**
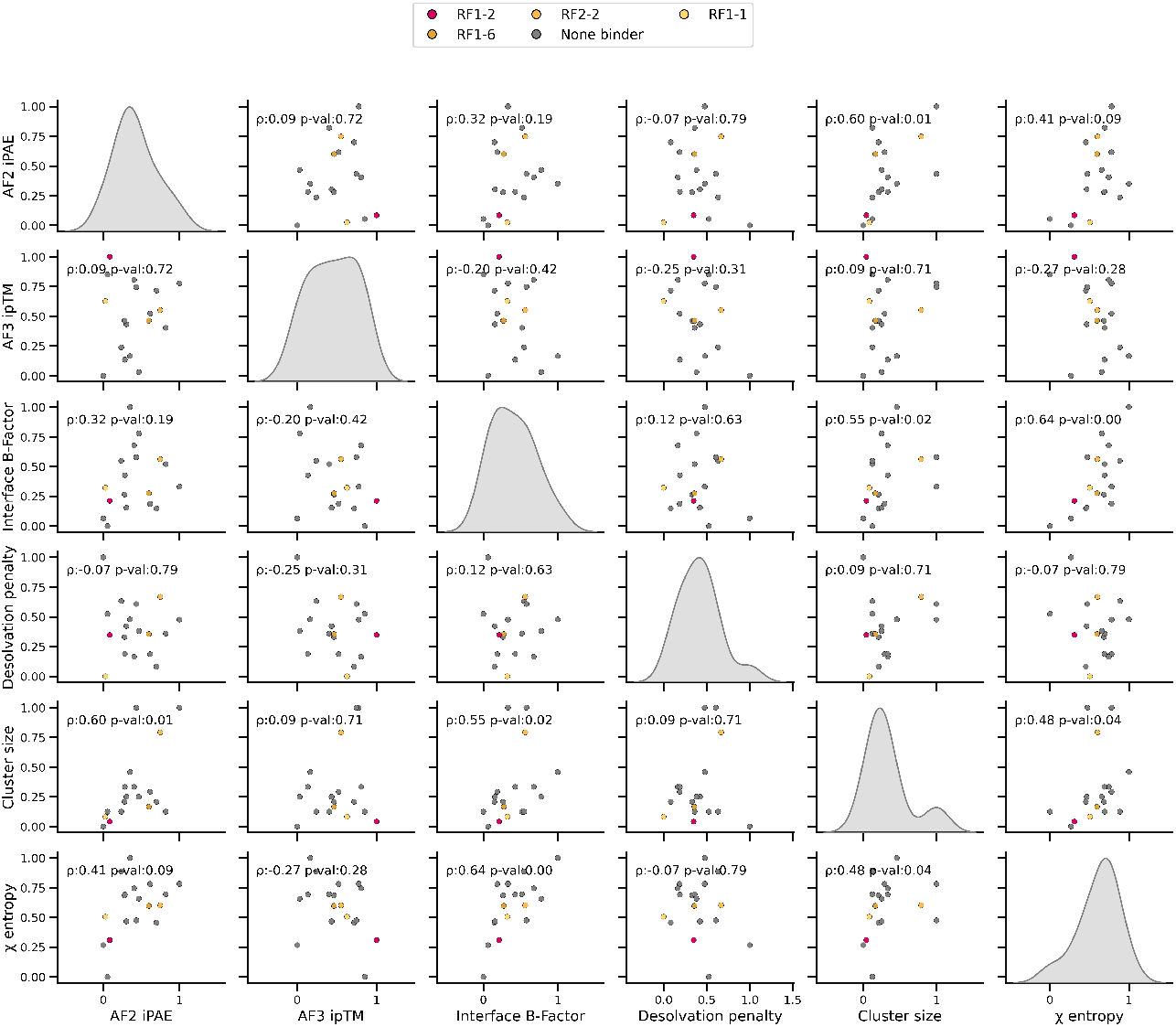
Pairwise scatter plots of min–max normalized MD features and static cofolding metrics, with Spearman rank correlations (*ρ*) annotated.

**Table 1:** MiBds selected for *in vitro* validation with final AF2-*ig* metrics, designed sequence length, and AF2-*ig* iPAE of initial binder designs.

| Minibinder | iPAE [Å] | Binder pLDDT | Sequence length | Initial iPAE [Å] |
| --- | --- | --- | --- | --- |
| BETA1 | 6.963 | 90.382 | 104 | 13.312 |
| BETA2 | 8.212 | 91.148 | 124 | 13.217 |
| RF1-1 | 5.798 | 90.409 | 102 | 17.410 |
| RF1-2 | 6.010 | 93.493 | 128 | 19.418 |
| RF1-3 | 6.550 | 92.981 | 111 | 11.162 |
| RF1-4 | 6.718 | 92.092 | 101 | 18.184 |
| RF1-5 | 7.157 | 90.421 | 101 | 13.891 |
| RF1-6 | 7.853 | 90.628 | 111 | 18.258 |
| RF1-7 | 7.910 | 90.262 | 114 | 17.320 |
| RF1-8 | 8.646 | 91.128 | 134 | 15.751 |
| RF1-HS-1 | 5.705 | 93.462 | 102 | 7.375 |
| RF1-HS-2 | 5.900 | 93.201 | 115 | 8.480 |
| RF1-HS-3 | 6.704 | 92.579 | 136 | 9.711 |
| RF1-HS-4 | 6.791 | 92.028 | 102 | 9.050 |
| RF1-HS-5 | 7.375 | 93.600 | 134 | 8.151 |
| RF2-1 | 7.258 | 91.948 | 144 | 9.752 |
| RF2-2 | 8.382 | 89.366 | 146 | 10.370 |
| RF2-3 | 9.288 | 91.847 | 143 | 12.241 |

**Table 2:** Descriptor-values derived from molecular dynamics analysis.

| Minibinder | Interface B-Factor | Desolvation | Cluster (aligned MB) | Chi entropy |
| --- | --- | --- | --- | --- |
| BETA1 | 155.10 | 99.80 | 12 | 3.71 |
| BETA2 | 60.85 | 63.16 | 6 | 1.98 |
| RF1-1 | 80.10 | 55.69 | 3 | 2.14 |
| RF1-2 | 67.87 | 87.78 | 2 | 1.52 |
| RF1-3 | 105.05 | 114.01 | 4 | 3.35 |
| RF1-4 | 91.77 | 73.10 | 9 | 2.73 |
| RF1-5 | 119.36 | 70.85 | 9 | 2.90 |
| RF1-6 | 74.98 | 88.51 | 5 | 2.43 |
| RF1-7 | 65.09 | 72.88 | 8 | 3.01 |
| RF1-8 | 101.99 | 88.85 | 4 | 2.74 |
| RF1-HS-1 | 51.56 | 147.94 | 1 | 1.39 |
| RF1-HS-2 | 44.55 | 104.07 | 4 | 0.54 |
| RF1-HS-3 | 73.79 | 86.13 | 6 | 2.71 |
| RF1-HS-4 | 61.59 | 94.55 | 7 | 2.02 |
| RF1-HS-5 | 130.51 | 90.83 | 7 | 2.62 |
| RF2-1 | 108.49 | 111.72 | 25 | 2.04 |
| RF2-2 | 106.64 | 117.25 | 20 | 2.45 |
| RF2-3 | 81.05 | 99.64 | 25 | 3.01 |

